# Temporal dynamics of microbiome communities within urban compost piles undergoing the heat process

**DOI:** 10.64898/2026.02.10.705224

**Authors:** Angel Montes, Dayton Klopmanbaerselman, Bertram Lee, Beatriz Quiñones, Hyunjin Shim

**Author notes:** Corresponding author: Hyunjin Shim.

## Abstract

Urban composting supports soil health but also intersects with food safety, where compost is produced near farms and communities. Here, we profiled temporal microbiome dynamics across a 6-week heat compost cycle from the urban compost piles using paired physicochemical panels and long-read metagenomics. Nutrient composition and pH shifted with compost age, coinciding with stage-structured microbial succession, including temperature-linked turnover of compost communities from mesophilic to thermotolerant taxa. Bacterial profiles included the presence of antimicrobial resistance genes and foodborne-associated genera early in the cycle, with reduced representation during the thermophilic phase. Analysis of previously unclassified long reads reveals an extensive repertoire of putative bacteriophages, including several complete genomes and candidates linked to foodborne bacteria, and their abundance is coupled to the host abundance. Together, these results support thermophilic composting as a key mitigation step for microbiological hazards in urban-adjacent systems and identify compost piles as a promising reservoir for discovering candidate lytic phages for downstream isolation and host-range testing.

## Introduction

Composting is a self-heating, biodegradable process that transforms organic materials into nutrient-rich soil through microbial decomposition (1). Successful composting depends on managing oxygen, moisture, carbon-to-nitrogen balance, pH, and temperature, with macronutrients (Nitrogen N, Phosphorus P, Potassium K) and other physicochemical properties changing as decomposition proceeds (2,3). Urban composting further intersects with public and occupational health because bioaerosols and residual microbial constituents may be encountered by workers and nearby communities. Understanding the microbiology of compost - what organisms are present, how they change over time, and how they relate to process conditions - is therefore central to both product quality and safe practice.

The heat (thermo-oxidative) phase of composting is a hallmark of this process and reflects the balance between microbial metabolism and heat loss. Classical composting proceeds through mesophilic, thermophilic, and maturation stages. During the mesophilic stage (20-40 ℃), fast-growing heterotrophs rapidly catabolize simple substrates, driving the temperatures upward (4,5). The thermophilic stage (40-70 ℃) follows, dominated by thermophilic bacteria, actinomycetes, and fungi that attack lignocellulose and other complex polymers; under appropriate time-temperature profiles, this phase also activates many plant pathogens and weed seeds (6–8). As easily oxidizable carbon is depleted and heat generation declines, piles cool and the community shifts back toward mesophiles, often yielding the most taxonomically diverse microbes. The maturation stage consolidates stability, further transforming recalcitrant substrates while pH, moisture, and nutrient forms approach their final values (9).

Previous studies have characterized microbial succession and physicochemical trajectories in compost using culture-based assays, molecular markers, and short-read metagenomics (8,10,11). These studies link temperature, moisture, pH, and nutrient availability to shifts in bacterial and fungal communities, and some have evaluated the fate of specific foodborne pathogens under controlled conditions (e.g. *Listeria monocytogenes*) (11). However, comparatively little attention has been paid to bacteriophages (phages) - the viruses of bacteria - which shape bacterial mortality, horizontal gene transfer, and metabolic potential across ecosystems. Phage lifestyles (virulent vs. temperate), host range, and gene cargo can modulate bacterial turnover and trait dissemination, but their roles in compost remain poorly resolved (12). This gap persists in part because fragmented assemblies and short reads complicate phage detection, host linkage, and lifestyle inference in complex, mixed communities (13,14).

Here, we address this gap by coupling time-resolved sampling of urban compost piles undergoing the heat process with long-read metagenomics and complementary chemical analyses. Long reads improve recovery of high-contiguity genomes, prophage boundaries, and mobile elements, enabling more confident discrimination of phage genomes and their bacterial hosts than short-read approaches alone (15,16). Composting underpins soil health and nutrient sustainability in California’s Central Valley, a major agricultural region. Our objectives were to: (i) quantify temporal changes in microbial community composition across the mesophilic, thermophilic, and maturation stages; (ii) relate microbial dynamics to temperature, pH, moisture, and macronutrient profiles measured over the same interval; and (iii) characterize phage diversity, predicted lifestyles, and putative host associations. Given the proximity of urban composting to population centers and food production, we also screened for bacterial taxa of health relevance, recognizing that metagenomic detection does not imply organism viability or activity.

From a medical and food-safety perspective, urban compost piles may function as reservoirs of novel bacteriophages with activity against priority foodborne bacteria. By applying long-read metagenomics with phage-centric prediction (lifestyle, host linkage, genome features), we interrogate the compost virome across the heat process to identify virulent (lytic) phages and track how phage presence aligns with the abundance of putative pathogens. By linking virome features to community stability and pathogen attenuation, our study positions compost as an accessible source of phages predicted to target bacterial pathogens and reaffirms the role of heat process in keeping communities safe while maintaining the microbial functions for efficient composting in urban-agricultural interfaces.

## Results

### Density of urban farming in Central California

Using state-curated aerial photography and field verification (17), we mapped agricultural land use across Fresno County (Figure 1A). Vineyards and deciduous orchards dominated the landscape: 6116 vineyard parcels accounted for 19.3% of total agricultural acreage, and 6202 deciduous orchard parcels accounted for 13.7%. Other major types of farming included field crops, citrus/subtropical, semi-agricultural, pasture, truck, nursery, and berry crops, underscoring the diversity of crop and agricultural production in Fresno County. Variations of parcel counts demonstrated the natural tendencies of agriculture in Fresno County, while the percentages of acreage revealed the sustainably dominant selection of high-value crops. These distributions emphasized the close spatial coupling of intensive agriculture with urban areas (Figure S1).

**Figure 1:**
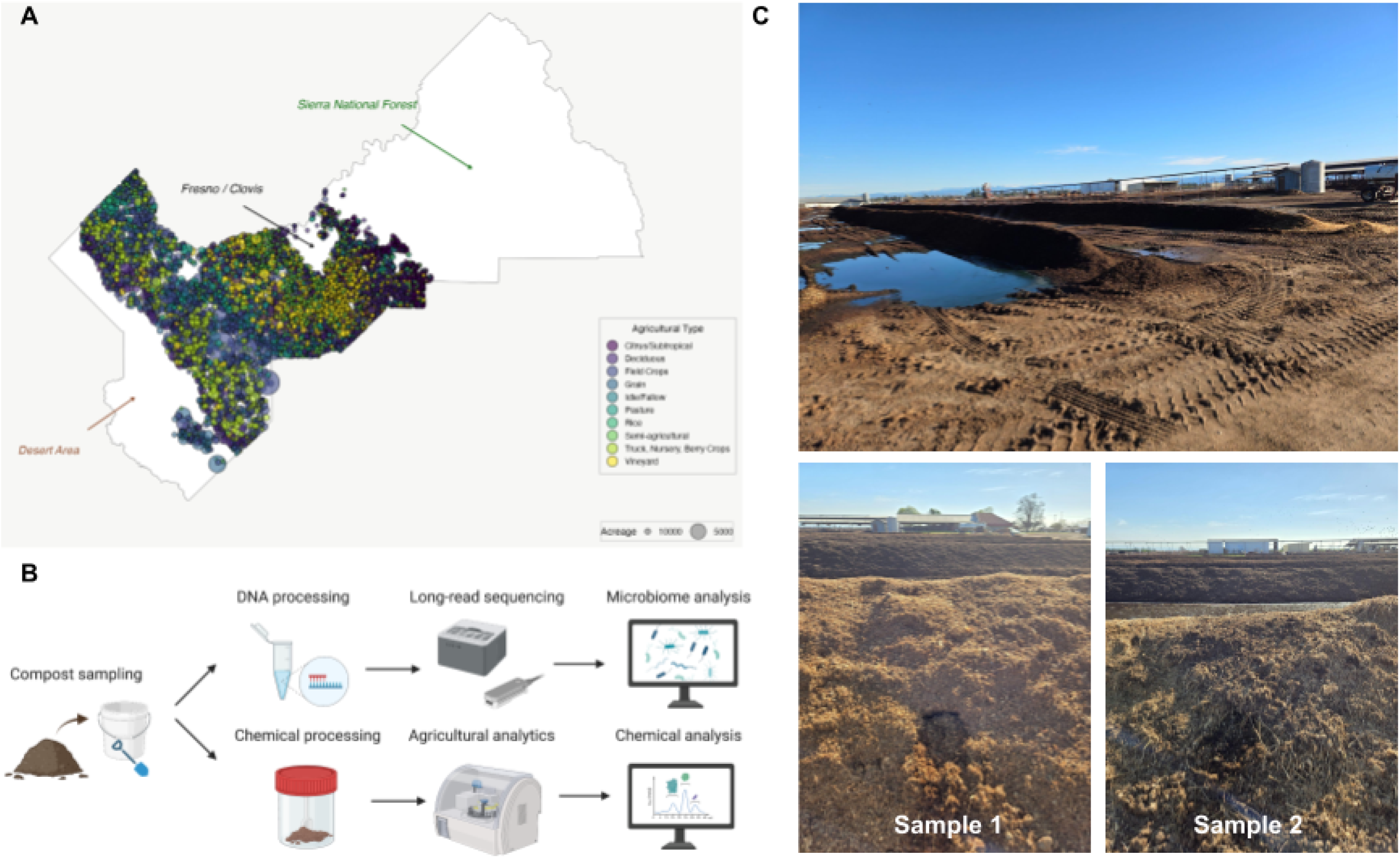
Study site and field sampling for urban compost heat-cycle metagenomics. (A) Agricultural land-use intensity in Fresno County (50). Circles denote individual agricultural parcels from a randomly selected 50% subset of field records; circle size scales with parcel acreage, and color indicates crop category. Major geographic landmarks (Sierra National Forest, Fresno/Clovis metropolitan area, and adjacent desert region) are labeled for orientation. The 50% subset is shown to reduce overplotting and improve visualization of crop-type diversity relative to the full dataset (Figure S1). (B) Overview of sampling-to-analysis workflow. Compost material collected from each pile was homogenized and split into two aliquots: one for long-read DNA extraction and sequencing performed in-house, and one for physicochemical characterization performed by a local agricultural consulting laboratory (Pacific Agronomics, USA). (C) Urban compost piles undergoing the heat process at the California State University, Fresno farm unit. Two independent biological replicates (“Sample 1” and “Sample 2”) were collected from two separate piles at each time point. For each collection, a new side of the pile was exposed, and material was sampled aseptically from the active thermophilic zone.

Our study site was located within this matrix (Figure 1C). California State University, Fresno, centrally located in Fresno County, operates ∼1000 acres across 20 farm units that collectively simulate urban-adjacent agriculture, including an active heat-compost field (18). The University Agricultural Laboratory, Animal Science facilities, and research units contribute to the addition of many crops, such as almond orchards and the breeding of livestock (18), including ∼120 acres of table, raisin, and wine grape vineyards; ∼325 acres of orchards (e.g. almonds, grapefruit, mandarins, nectarines, olives, oranges, peaches); and ∼200 acres of both vegetable and field crops (e.g. corn).

### Temporal changes in compost physicochemical properties

We quantified macronutrients and key physicochemical parameters over the compost heat cycle, with two independent biological replicates per time point (Figure S2; Table S1). Because Sample 2 began in an advanced state at Week 1, we present Sample 1 (new pile) as the primary trajectory and analyze both samples together after aligning by compost age (Figure 2). Nitrogen content exhibited the expected rise-fall pattern (19): it increased from 1.08% at initiation to 1.33%, then declined to 0.58% by the end of the cycle (Figure 2A). Phosphorus showed modest variation as expected (20), increasing from 0.24% to 0.39%, before returning to 0.19% (Figure 2B). Potassium decreased from 1.61% to 0.62% overall (Figure 2C), indicating a net potassium loss with leaching or biomass incorporation despite the commonly modest K fluctuation reported in other systems (20). The amount of zinc in a pile should relatively remain constant, but the concentration of the nutrient in ppm should increase throughout the composting process (21). Zinc concentrations increased mid-cycle to 58 ppm and declined to 33 ppm at the end (Figure 2D). A compost pile’s pH levels should begin at a slightly acidic range and then increase to slightly alkaline before neutralizing back to around a pH level of about 6.5-7.5 (19). The pH values followed the canonical trajectory of starting at 6.8, peaking at 8.1 (Week 3), dropping to 7.5 (Week 5), then rising again towards the end of the heat cycle (Figure 2E).

**Figure 2:**
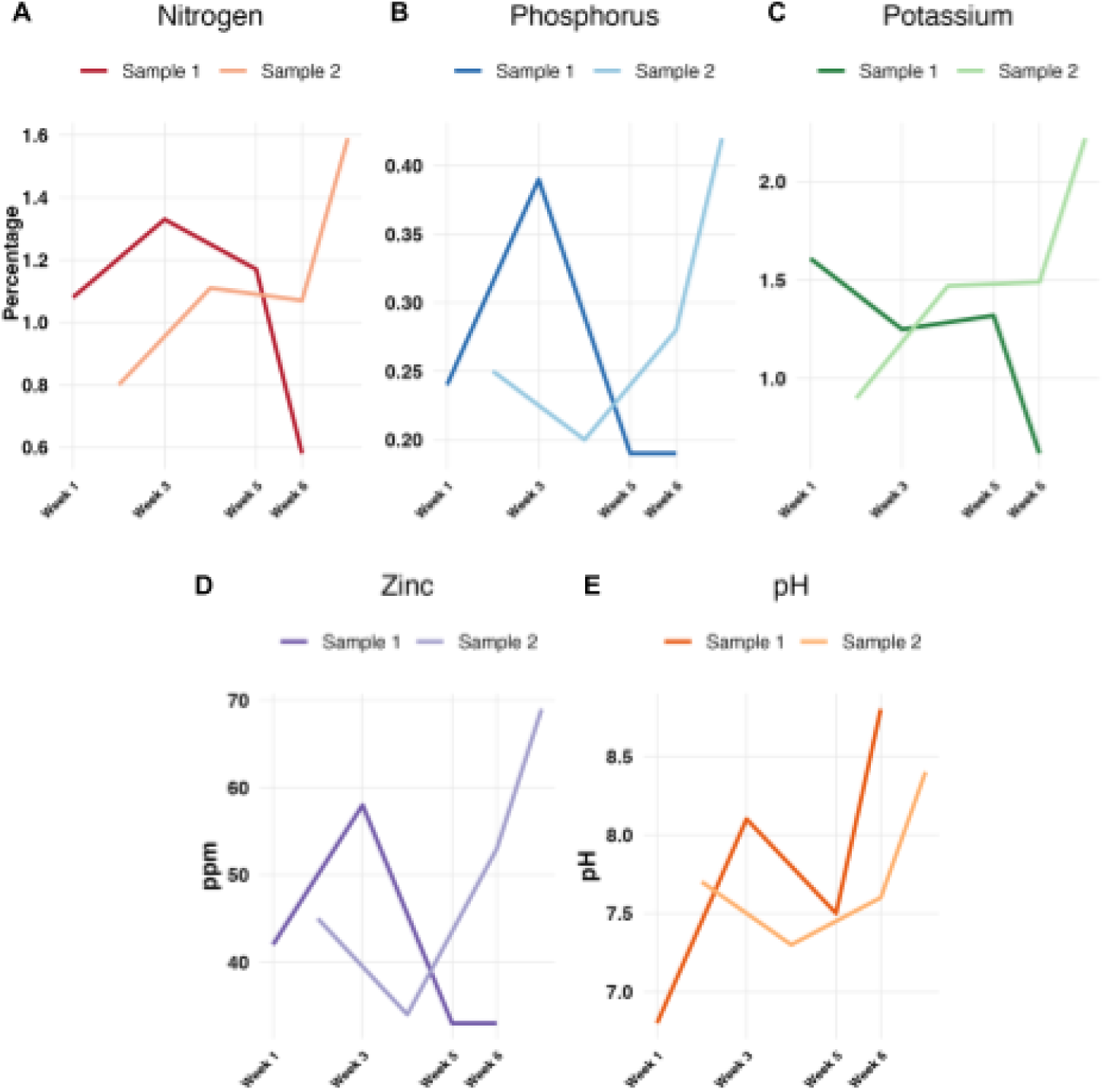
Temporal changes in compost physicochemical properties across the heat process. Chemical profiles were measured for two independent compost piles (Sample 1 and Sample 2) across the heat cycle. Sample 2 time points are shown shifted by one week to align the compost age with Sample 1 (see Figure S2). (A) Total nitrogen (%); (B) Total phosphorus (%); (C) Potassium (%); (D) Zinc (ppm); (E) pH.

After shifting Sample 2 by one week to match the compost age with Sample 1 (Figure 2), replicate trends converged. Nitrogen increased from a mean of 0.94% (Weeks 1-2) to 1.22% at Week 3, then declined to 1.085% at the end of the cycle (median of 0.58% to 1.59%, average SD 0.31%) (Figure 2A). Phosphorus remained relatively stable at 0.245-0.295% during the first four weeks before rising to 0.305% at the end of the cycle (median of 0.19 to 0.42%, average SD 0.09%) (Figure 2B). Potassium remained within the ranges of 1.26% to 1.36% for the first four weeks and increased to 1.42% at the end of the cycle (median of 0.62% to 2.22%, average SD 0.48%) (Figure 2C). Zinc increased from a mean of 43.5 ppm (Weeks 1-2) to 51 ppm at the end of the cycle, with growing dispersion (median of 33 ppm to 69 ppm, average SD 13.2 ppm) (Figure 2D). Finally, pH progressed from a mean of 7.25 (Weeks 1-2) to 8.60 by the end of the cycle (median of 8.4 to 8.8, average SD 0.64) (Figure 2E).

### Temporal changes in compost microbiome compositions

The soil compositions were monitored with long-read sequencing for the 6-week compost cycle (Tables 1, S2). The nanopore output showed that soil compositions were dominated by bacteria, while up to 70% of the reads remained unclassified from the Kraken taxonomic classification database (Table S3) (22). These compositions fluctuated in time; the bacterial compositions tended to increase in the middle of the cycle, before decreasing towards the end of the cycle (Figure S3). The other superkingdoms (archaea, eukaryotes, viruses) were observed to have slight changes over time but maintained a prevalence under 1.5% for eukaryotes, 0.5% for archaea, and 0.1% for viruses throughout the compost cycle.

**Table 1:** Sampling schedule and Nanopore sequencing yields from a 6-week urban compost heat cycle. Two independent biological replicates were collected at each time point from two separate compost piles (Sample 1 and Sample 2). Sequencing yield is reported as total base-called output in gigabase pairs (Gbp) per sample.

|  |  | Sample 1 |  |  |  | Sample 2 |  |  |  |
| --- | --- | --- | --- | --- | --- | --- | --- | --- | --- |
| Week | Date | DNA quantity<br>(ng/μl) | Data output<br>(Gb) | Read count<br>(M) | N50<br>(kb) | DNA quantity<br>(ng/μl) | Data output<br>(Gb) | Read count<br>(M) | N50<br>(kb) |
| 1 | 2024-03-11 | 2480 | 26.79 | 9.51 | 3.60 | 2347 | 12.95 | 4.37 | 4.72 |
| 3 | 2024-03-25 | 2252 | 25.22 | 7.97 | 4.63 | 1817 | 16.72 | 9.15 | 2.75 |
| 5 | 2024-04-08 | 1965 | 12.83 | 5.07 | 3.61 | 1682 | 15.47 | 5.52 | 4.47 |
| 6 | 2024-04-15 | 1612 | 17.17 | 8.87 | 2.96 | 1598 | 24.30 | 6.85 | 4.94 |

The most abundant phyla during the compost process were from bacteria, such as Pseudomonadota, Actinomycetota, Bacillota, Bacteroidota, Thermomicrobiota, and Planctomycetota (Table S3). Examples of species within these phyla included *Escherichia coli* and *Pseudoxanthomonas suwonensis* for Pseudomonadota, *Thermobifida fusca* and *Mycolicibacterium thermoresistibile* for Actinomycetota, and *Sphaerobacter thermophilus* for Thermomicrobiota (Figure 3A-B). Among the top ten phyla, the results also identified *Chordata* and *Streptophyta* from Eukaryota, and *Euryarchaeota* from Archaea (Figure S4). The most abundant phyla that remained relatively stable over the compost cycle, except for the Sample 2 last sampling point included Uroviricota from Viruses (Table S3). Uroviricota is a class of *Caudoviricetes*, which are tailed and head-tail viruses with double-stranded DNA genomes (23). Given that the last sampling point for Sample 2 represented a new pile undergoing the initial compost cycle, the role of *Caudoviricetes* in the compost cycle was found to be notable. In particular, most *Caudoviricetes* belonged to a family of *Herelleviridae* (70% of *Caudoviricetes* reads), infecting members of the phylum Bacillota (24), which was the third most abundant phylum at this time point of Sample 2.

**Figure 3:**
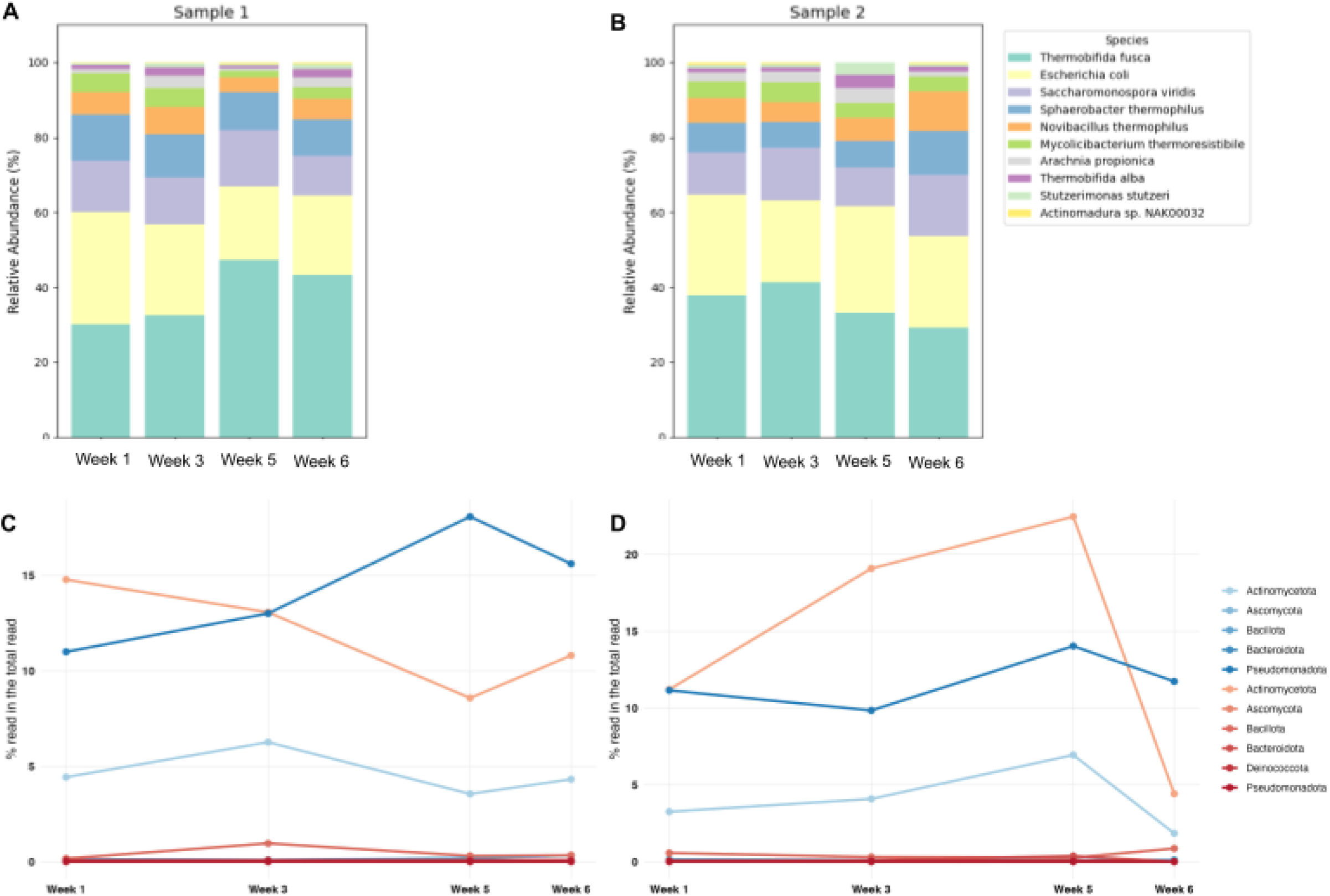
Temporal changes in compost microbiome composition across the heat process. Taxonomic profiles were generated from quality-filtered Nanopore reads using a minimum read length of 200 bp. (A-B) Relative abundance of the top ten bacterial species in Sample 1 (A) and Sample 2 (B), stacked bars show the fraction of total classified reads per time point. (C-D) Temperature-associated bacterial community structure in Sample 1(C) and Sample 2 (D), summarized at the phylum level and grouped by thermal preference. Taxa classified as mesophilic are shown in blue, and taxa classified as thermophilic are shown in red.

During the mesophilic interval (Weeks 1-3), communities were enriched for mesophilic taxa under moderate temperatures, consistent with early substrate turnover (Figure 3C). Mesophilic taxa, including members of Ascomycota, were relatively abundant and initiated degradation of readily available substrates (e.g. simple carbohydrates), consistent with rapid primary decomposition. As the pile transitioned to thermophilic conditions (Weeks 4-5), heat-tolerant fungi increased in relative abundance, with Ascomycota remaining prominent, indicating a shift towards thermophilic fungi capable of withstanding elevated temperatures and attacking more complex polymers (25). This temperature-driven succession, in which different members of the same phylum contribute at distinct stages, is well documented for composting systems and supports a mesophile-to-thermophile turnover of functional fungi (26). Overall, this observed shift from mesophilic to thermophilic fungi species matches canonical compost succession and underscores stage-specific selection of fungal decomposers across the heat process.

### Detection of pathogenic microbes in the compost piles

As bacterial constituents of compost can influence human and environmental health, we performed a bacteria-centric analysis of the metagenomic reads. The most abundant bacterial species included the thermophile *Thermobifida fusca* and the mesophiles *E. coli* throughout the compost cycle, except for the last sampling time point of Sample 2, with increased abundance of other taxa (Table S4). Mesophilic bacteria detected included *Escherichia coli, Salmonella entericae,* and *Pseudomonas aeruginosa*, while the thermophilic bacteria included *Thermobifida fusca, Thermobifida alba, Mycolicibacterium thermoresistibile, Actinomadura* sp. NAK00032, and *Actinomadura madurae* (Table S5). Notably, reads specific for the foodborne pathogen *S. enterica* and generic *E. coli* were food-relevant pathogens; their reads were most evident during cooler phases and declined as piles reached thermophilic temperatures, consistent with temperature-driven reduction of pathogens during composting (26).

Based on the foodborne pathogen results analysis, a high proportion of *Salmonella*-specific reads were identified in both Sample 1 and 2 for all examined dates (Figure 4), and *Salmonella* was still the most common foodborne pathogenic species detected during week 5 for Sample 1. Interestingly, the relative abundance of reads, corresponding to the bacterial pathogens *Brucella* spp., *Bacillus cereus*, *Listeria monocytogenes*, *Staphylococcus aureus*, and *Vibrio* spp. were observed to increase in the last sampling date in Week 6 for Sample 2, which was previously identified as the start of a new pile. The relative prevalence of *Bacillus cereus* and *Vibrio* spp. increased 4 to 5-fold, and notably, the prevalence of *Listeria monocytogenes* and *Staphylococcus aureus*, pathogens implicated in causing severe human illness, increased over 10-fold (Table S6).

**Figure 4:**
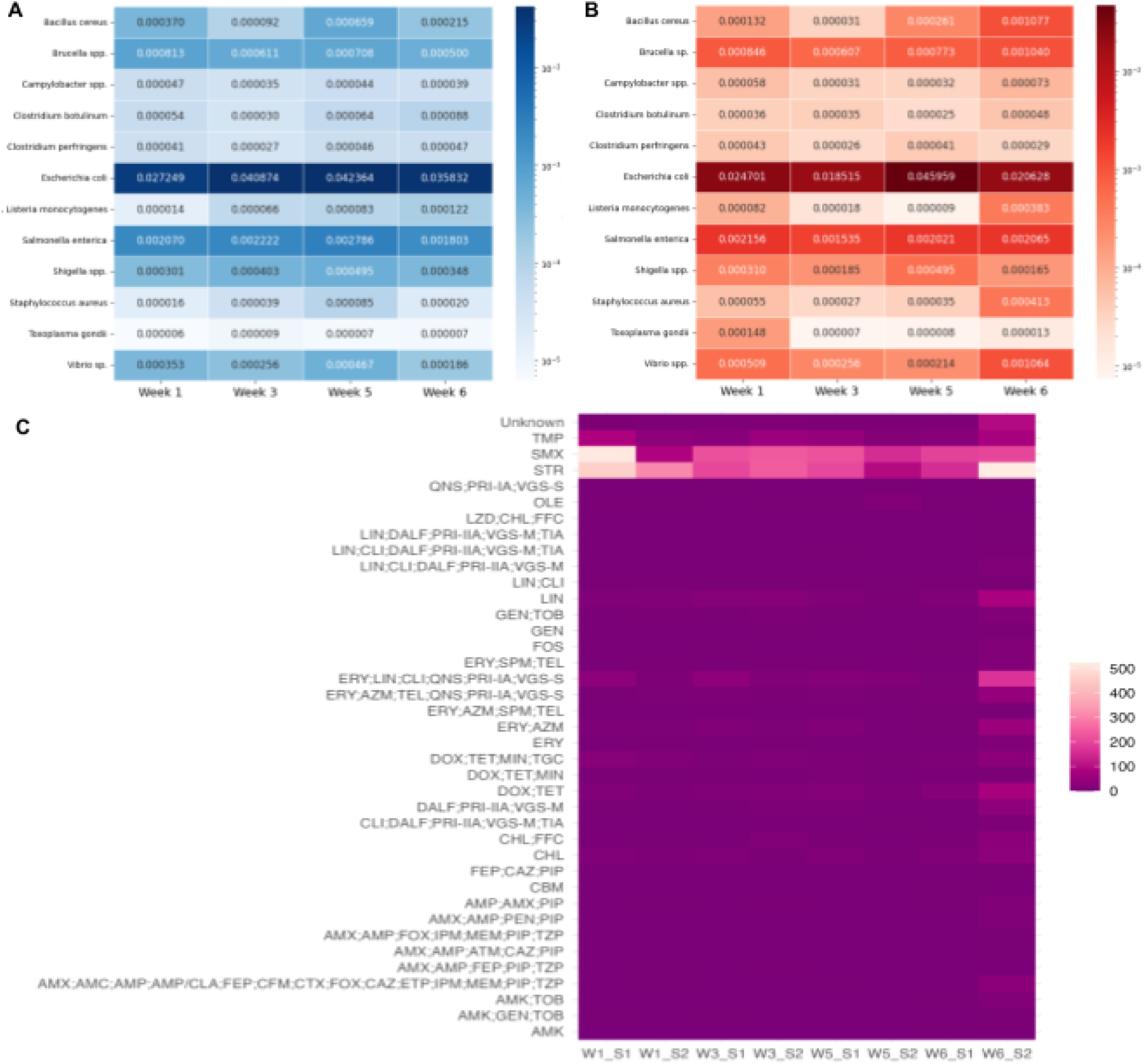
Temporal dynamics of pathogenic bacteria and antimicrobial resistance across the compost heat process. (A-B) Heatmap of the proportion of the reads predicted to be foodborne bacteria across the sampling time point in Sample 1 (A) and Sample 2 (B). (C) Heatmap of antimicrobial resistance (AMR) genes detected from metagenomic contigs using Abricate with the ResFinder database. Cell values represent the number of AMR gene hits per sample/time point. Antibiotics (Abbreviation) is given as Amikacin (AMK); Gentamicin (GEN); Tobramycin (TOB); Amoxicillin (AMX); Amoxicillin + Clavulanic acid (AMC); Ampicillin (AMP); Ampicillin + Clavulanic acid (AMP/CLA); Piperacillin (PIP); Piperacillin + Tazobactam (TZP); Cefepime (FEP); Cefixime (CFM); Cefotaxime (CTX); Cefoxitin (FOX); Ceftazidime (CAZ); Ertapenem (ETP); Imipenem (IPM); Meropenem (MEM); Aztreonam (ATM); Penicillin (PEN); Carbomycin (CBM); Chloramphenicol (CHL); Florfenicol (FFC); Clindamycin (CLI); Dalfopristin (DALF); Pristinamycin IIA (PRI-IIA); Virginiamycin M (VGS-M); Tiamulin (TIA); Doxycycline (DOX); Tetracycline (TET); Minocycline (MIN); Tigecycline (TGC); Erythromycin (ERY); Azithromycin (AZM); Spiramycin (SPM); Telithromycin (TEL); Quinupristin (QNS); Pristinamycin IA (PRI-IA); Virginiamycin S (VGS-S); Linezolid (LZD); Oleandomycin (OLE); Streptomycin (STR); Fosfomycin (FOS); Sulfamethoxazole (SMX); Trimethoprim (TMP); Lincomycin (LIN).

We also detected numerous antimicrobial resistance (AMR) genes across samples (Table S7). Counts were highest for genes conferring resistance against trimethoprim (TMP), sulfamethoxazole (SMX), and streptomycin (STR), each exceeding 200 hits at one or more time points, whereas determinants conferring resistance to other agents (e.g. beta-lactams, carbapenems, tetracyclines, phenicols, fosfomycin, aminoglycosides) generally remained below 200 hits (Figure 4C). A subset of loci could not be unambiguously assigned to a single drug class (“unknown”). Notably, Sample 2 at Week 6 (the start of a new pile) showed a broad uptick in gene counts, including over 200 hits spanning the macrolide-lincosamide-streptogramin (MLS) group, consistent with an early-cycle enrichment of resistance loci that subsequently declines as composting progresses. Together with the foodborne-pathogen screen, the patterns observed in the metagenomic analyses suggest that thermophilic composting can potentially reduce some risks related to pathogen and AMR gene prevalence over time, likely via elevated temperature, aeration, and competitive dynamics during active decomposition (30,31).

### Discovery and characterization of novel bacteriophages in the compost piles

After applying a length filter of 3,000 bp, a substantial fraction of previously unclassified Nanopore reads were predicted as putative phages (Table S8). In Sample 1, phage-assigned reads comprised 0.54-1.5% of totals, peaking at 1,585,813 reads in Week 1. For Sample 2, phage-assigned reads comprised 0.58-5.4%, peaking at 1,253,424 reads in Week 6 (the start of a new heat cycle). These observations showed that compost piles harbor an untapped repertoire of unclassified phages, which are particularly prominent at the beginning of a heat process. Length distributions were broad, with a median length near 4 kb and maxima reaching around 50 kb at all time points (Figure 5A), consistent with near-complete tailed-phage genomes captured as single long reads (32). These results indicate that long-read assemblies of compost materials yield high-contiguity viral sequences suitable for downstream analyses of phage lifestyle, host association, and therapeutic candidate selection.

**Figure 5:**
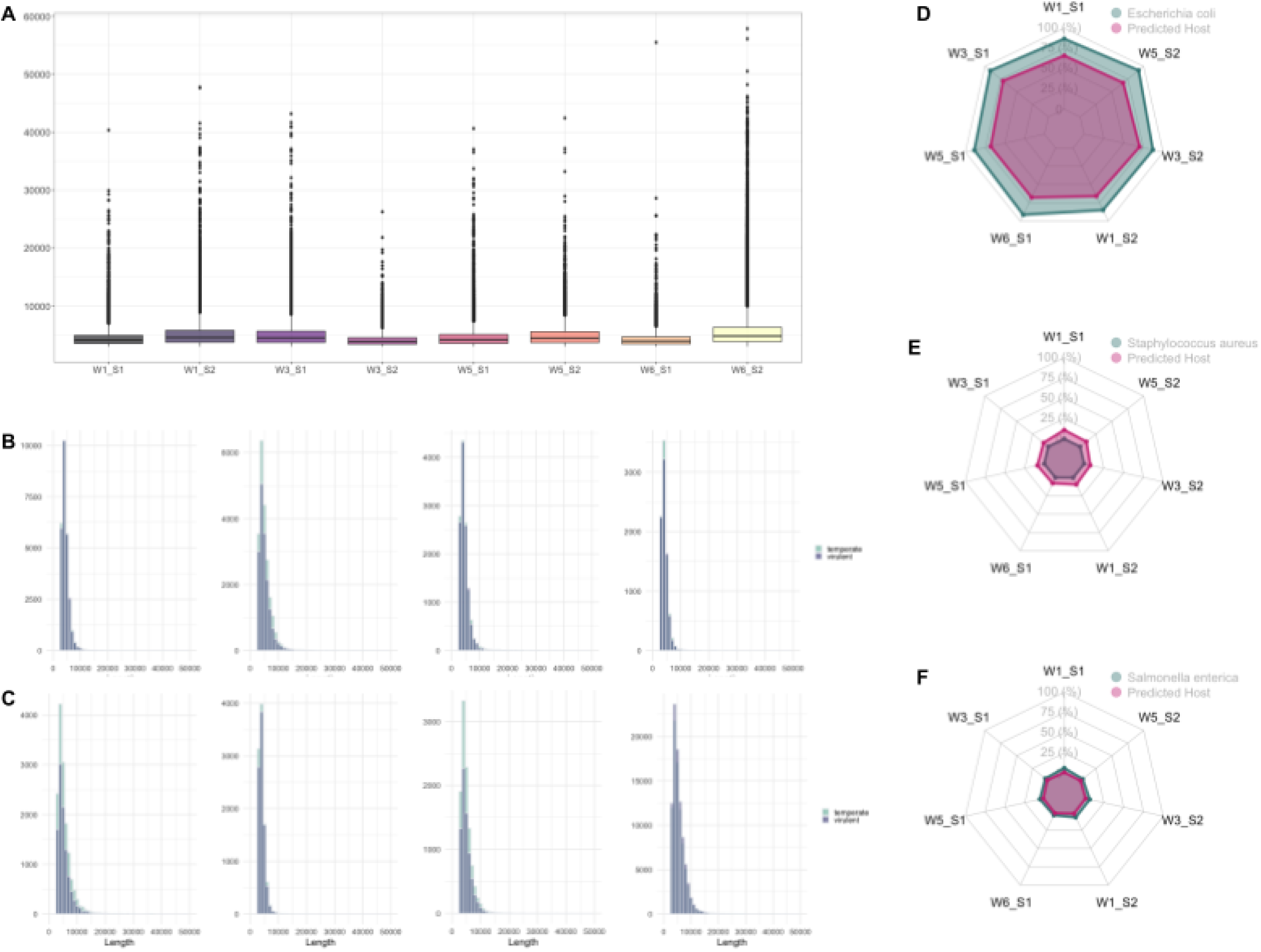
Characterization of putative bacteriophages from previously unclassified reads. The identification of phage sequences from reads not initially assigned by the taxonomic profiling was performed by further screening with a minimum length threshold of 3,000 bp prior to prediction. (A) Length distributions of unclassified reads predicted as “virus” by PhaBOX, highlighting recovery of long fragments consistent with high-contiguity phage genomes. (B-C) Length distributions of predicted phage sequences stratified by lifestyle (virulent or temperate) for Sample 1 (B) and Sample 2 (C). (D) Radar plots comparing the relative abundance of foodborne-associated bacterial taxa and the corresponding abundance of phage contigs with predicted hosts matching those taxa to illustrate phage-host coupling during the compost heat process.

Within each sample, genome-length distributions did not differ significantly between reads predicted as virulent (lytic) versus temperate (lysogenic) (Figure 5B-C), indicating that genome size alone is not a good predictor for phage lifestyle in this system (33). Nevertheless, virulent predictions highlighted promising biocontrol candidates for isolation and host-range testing. Host-prediction summaries (Table 2) aligned with the foodborne bacterial species detected (Figure 4), yielding multiple candidate phages against pathogen targets of interest (e.g. *Listeria* spp., *Salmonella* spp., Shiga toxin-producing *E. coli*, *Campylobacter* spp.), whereas only single-digit candidates were recovered for *Clostridium botulinum* and *Clostridium perfringens*. This likely reflected low host abundance in our dataset (Table S6) and/or limited representation of phages associated with these bacterial pathogens in the training database for the machine learning method (34). The number of predicted phage hosts per taxon correlated with the relative abundance of corresponding bacteria in the radar chart (Figure 5C), suggesting phage-bacteria coupling in these communities. The observed pattern in the present study was found to be consistent with previous studies examining phage-mediated modulation of bacterial populations during the compost heat process (13,35,36).

**Table 2:** Putative phage contigs from previously unclassified reads with predicted hosts matching foodborne-associated bacteria. Unclassified Nanopore sequences were screened, and those predicted as phages were linked to candidate bacterial hosts using PhaBOX host prediction. Assigned host: a curated set of foodborne-associated taxa used in this study Phage contigs: the number of unclassified reads predicted to be phage-derived for each host

|  | Sample 1 |  |  |  | Sample 2 |  |  |
| --- | --- | --- | --- | --- | --- | --- | --- |
| Assigned host | W1_S1 | W3_S1 | W5_S1 | W6_S1 | W1_S2 | W3_S2 | W5_S2 |
| *Escherichia coli* | 2304 | 1747 | 1245 | 985 | 1077 | 1085 | 749 |
| *Salmonella enterica* | 44 | 43 | 28 | 25 | 41 | 31 | 7 |
| *Shigella spp.* | 26 | 15 | 13 | 14 | 9 | 18 | 11 |
| *Bacillus cereus* | 153 | 119 | 62 | 100 | 121 | 73 | 100 |
| *Brucella sp.* | 94 | 48 | 56 | 37 | 28 | 40 | 20 |
| *Campylobacter sp.* | 46 | 22 | 25 | 22 | 15 | 20 | 8 |
| *Clostridium botulinum* | 0 | 0 | 0 | 0 | 1 | 0 | 0 |
| *Clostridium perfringens* | 1 | 6 | 1 | 0 | 2 | 3 | 0 |
| *Listeria monocytogenes* | 113 | 93 | 58 | 63 | 91 | 50 | 39 |
| *Staphylococcus aureus* | 387 | 185 | 163 | 113 | 154 | 122 | 110 |
| *Toxoplasma gondii* | 0 | 0 | 0 | 0 | 0 | 0 | 0 |
| *Vibrio spp.* | 311 | 169 | 182 | 95 | 95 | 104 | 63 |

A noteworthy exception is the low number of phage contigs predicted for *Salmonella enterica* (Table 2), despite its high prevalence among pathogen-assigned reads. This discrepancy may reflect host-range gaps in current phage models, strain-specific or plasmid-mediated signals that confound host linkage, or differences in genome recoverability (e.g. fragmented Salmonella phage genomes below length thresholds). It may also mirror ecological differences in phage availability between *E. coli* and *Salmonella* in compost matrices. From the subset of viral predictions passing the length and confidence thresholds, we recovered several complete and near-complete phage genomes with predicted hosts linked to foodborne-associated bacteria, including two contigs with 94% and 100% genome completeness against *Salmonella entrica* (Table S9).

## Discussion

Urban composting is considered to be at the interface of sustainable agriculture, waste management, and public health, particularly in agricultural regions in California’s Central Valley, where intensive production systems depend on soil fertility. Within this context, California State University, Fresno’s farm units provide a representative, urban-adjacent platform to study compost microbiology. Our work advances this goal by integrating longitudinal and physicochemical profiling with long-read metagenomics, enabling simultaneous tracking of microbial succession and mobile components (including phages and resistance genes) across the compost heat process (37,38). The resulting framework is intended to support evidence-based compost management and to expand microbial ecology beyond bacteria/fungi to include phage-mediated regulation, which remains underrepresented in compost research (39).

Across the heat cycle, nutrient and microbial patterns were broadly consistent with canonical compost succession. After aligning the two piles by compost age, macronutrients, and pH, the findings followed coherent trajectories: nitrogen rose early and declined towards the end of the cycle, phosphorus remained comparatively stable with modest late increases, and pH shifted towards alkalinity during the thermophilic interval. These physicochemical changes coincided with clear community turnover, including temperature-driven dynamics among fungi. Several abundant bacteria were observed to have traits aligned with compost transformation. *P. aeruginosa* is a mesophile with documented capacities for degrading hydrocarbons and other organics, potentially accelerating mineralization of organic matter, reducing the localized anaerobic environment, and promoting further decomposition of organic matter for other microbes during the composting cycle (27). Among thermophiles, *T. fusca* is a known degrader of lignocellulose and other complex compounds, facilitating subsequent enzymatic steps by the broader community (28). *T. alba* is a heat-tolerant bacterium that can degrade aliphatic-aromatic copolyester film and decrease polymer particle sizes, highlighting complementary routes for complex-substrate turnover (29). Together, these mesophilic and thermophilic taxa illustrate a division of decomposition across the heat process, with early-stage mesophiles initiating substrate breakdown and thermophiles sustaining high-temperature degradation of complex organics. These results reinforce that composting is a staged ecological process driven by temperature, substrate availability, and changing chemistry that selects for different decomposer consortia over time (40).

A key motivation for profiling microbiomes in urban compost is the potential exposure to foodborne bacterial pathogens through direct handling, aerosolization, or downstream contact with crops. We detected sequences assigned to foodborne pathogens (e.g. *Escherichia* and *Salmonella*) predominantly during cooler phases, with reduced representation as piles reached thermophilic conditions. This pattern is consistent with the longest-established principle that well-managed heat composting reduces pathogen burden over time (26). Importantly, we emphasize two interpretive limits: metagenomic detection does not imply viability, and gene presence does not imply phenotypic resistance. Nonetheless, the convergence of early-cycle enrichment and later-cycle attenuation supports the practical value of achieving robust thermophilic conditions in reducing microbiological hazards, as discussed in recent compost-AMR studies highlighting the influence of temperature, aeration, moisture, and pH or AMR persistence (41,42).

A key contribution of this study is the recovery of a substantial “dark” virome from long reads that were initially unclassified by standard taxonomic workflows. A notable fraction of previously unclassified reads were predicted as phages, including sequences approaching ∼50 kb, consistent with near-complete tailed-phage genomes recovered at high contiguity. Lifestyle prediction indicated the presence of virulent candidates relevant for translational biocontrol due to the reduced concern for lysogeny and horizontal gene transfer (43,44). Host-prediction outputs further suggested that phage-host signals broadly tracked bacterial abundance patterns, consistent with phage-bacteria coupling, such as “kill-the-winner” dynamics that can modulate community structure during decomposition (45). Beyond ecological insight, the identification of candidate phages predicted to target foodborne bacteria positions compost piles as reservoirs of therapeutic phages targeting bacterial pathogens and phage-encoded antimicrobials. These observations align with the growing interest in phage-based approaches for managing bacterial infections (46,47).

This work also has several limitations, including the limited set of time points and sampling replicates. Expanding replication across seasons, feedstocks, turning regimes, and multiple independent compost sites will be necessary to generalize the observed trajectories. Furthermore, both phage lifestyle and host predictions are model-dependent; underrepresentation of certain hosts and uneven training data can bias host-linkage and partially explain underestimation, such as the genus of *Clostridium* (14). Finally, long-read metagenomics improves contiguity in the genomic information but does not address organism viability or activity. Future work will integrate pathogen viability and AMR assays, targeted phage isolation, and metatranscriptomics to distinguish live organisms and active phage infection cycles from residual DNA that persists in the environment. Despite these constraints, the combination of chemistry, longitudinal metagenomics, and long-read phage discovery provides a comprehensive study for evaluating how composting practice shapes microbial ecology while potentially impacting human health over time.

## Materials and Methods

### Temporal sampling of compost undergoing the heat process

Sampling was conducted at the California State University, Fresno farm unit over a single 6-week composting heat cycle (Figure 1B). To measure temporal dynamics, we sampled at four time points corresponding to early, mid, and late thermophilic phases (Table 1). At each time point, two independent biological replicates (“Sample 1” and “Sample 2”) were collected from two separate compost piles to quantify within-time-point variability.

On each date, a new side of the pile was exposed, and the weathered exterior layer was discarded to minimize exogenous contamination (Figure 1C). Using sterile sampling tools and nitrile gloves, material was collected aseptically from the active thermophilic zone of the pile. Each replicate was taken from separate piles to ensure independence while representing the same sampling time point. Samples were placed into pre-labeled sterile containers, sealed immediately, and tools were disinfected between replicates to prevent cross-contamination.

### Chemical analysis of soil samples from compost piles

All specimens were transported to the laboratory the same day, where they were homogenized and aliquoted for chemical composition analysis (Pacific Agronomics, USA) and long-read sequencing (Oxford Nanopore Technologies, UK).

For chemical analysis, ∼500 g of material per sample was placed in sterile, airtight containers and delivered to the analysis site. Chemical characterization was performed by a local agricultural consulting company (Pacific Agronomics, USA), using their standard compost analysis panel. Briefly, this panel typically reports moisture content, pH, electrical conductivity, total nitrogen, macronutrients (e.g. P, K), micronutrients (e.g. Zn), phosphate (P₂O₅), and potash (K₂O), by following common agronomic laboratory practices and methods. The laboratory provided QA/QC summaries and results as certified reports (Table S1).

### Nucleic acid extraction and ligation library preparation for metagenomic Nanopore Sequencing

Compost microbiome DNA was extracted from each sample using a soil-optimized kit (DNeasy PowerSoil Pro Kit; Qiagen, Germany), with bead beating and inhibitor removal, following the manufacturer’s instructions. Nucleic acids were purified with the kit’s recommended cleanup steps. DNA quantity and purity were assessed using a Qubit Fluorometer and Qubit dsDNA BR assay (ThermoFisher, USA), as summarized in Table 1.

Metagenomic libraries were prepared from each DNA extract using the ligation-based chemistry compatible with R10.4.1 flow cells and adhering to the supplier’s protocol (SQK-LSK114; Oxford Nanopore Technologies, UK). Briefly, high-integrity DNA was end-repaired/dA-tailed, followed by ligation of ONT sequencing adapters and post-ligation cleanups, as specified by the kit. Library concentration was verified using a NanoDrop microvolume spectrophotometer (ThermoFisher, USA).

Prior to loading, R10.4.1 flow cells were checked for pore availability (>800 active pores) using the ONT device control software, and were primed following the supplier’s protocol. Libraries were loaded to the MinION device (Mk1B; Oxford Nanopore Technologies, UK) and sequenced under standard MinKNOW (v23.11.2) run settings for ligation libraries. Run metrics (active pores, yield, read-length distributions) were monitored over a 72-hour acquisition. Resulting POD5 files were exported for downstream metagenomic analyses (Table S2).

### Downstream metagenomic analysis of Nanopore reads

Basecalling was performed with Dorado (v7.2.11) using the super-accuracy model appropriate for R10.4.1 chemistry. Reads were filtered to retain at a minimum per-read quality threshold of Qscore ≥ 10; reads failing this threshold were excluded from downstream analysis. Read-level quality control was summarized per sample using standard long-read QC tools of MinKNOW Core (v5.8.0). Metrics reported included total bases, read count, N50 read length, and passed/failed base calling counts (Table S1). The final high-quality FASTAQ files (Q ≥ 10) for each compost pile were carried forward to metagenomic analysis.

Taxonomic assignments were generated with the EPI2ME Labs workflow (v23.11.2) using Kraken2 with default parameters (22). We used the prebuilt PlusPFP-8 database with the index size of 7.5 Gb, which includes Archaea, Bacteria, viruses, plasmids, human, protozoa, fungi, plants (to capture plant-derived reads typical of compost feedstocks), and UniVec_Core to screen for vector contamination. Detailed comparisons throughout the compost process were created to analyze differences and trends between sampling points (Table S3-S5).

For species-level analysis, we applied a minimum detection threshold of 50 reads per species, based on a mock-community study indicating an elevated false positive rate below this level for Nanopore data (48). For the foodborne pathogen analysis, only species that had 50 reads in at least one time point were included in the analysis (Figures 4, S5). Moreover, the analysis of multiple species implicated in foodborne illness was performed by adding together the combined read counts for a particular genus, resulting in a combined read count higher than 50. In particular, the presence of the *Vibrio* genus with species only implicated in foodborne illness was considered in the analysis, even when the read counts were less than 50 for the individual species. Given that the genomic data did not identify virulence factors for the various *Escherichia coli* pathotypes, the results reported the prevalence of *Escherichia coli*, possibly encompassing pathogenic as well as non-pathogenic strains.

AMR genes were analyzed using Abricate (v1.0.1) with the ResFinder database. Reads were screened with a minimum identity of over 90% and coverage of over 80%. Hits corresponding solely to point-mutation mechanisms were excluded. For each sample, we report the AMR gene and the resistance drug for each sample (Table S7).

### Analysis of unclassified Nanopore reads

To mine the putative phage sequences missed by read-level taxonomy, the unclassified Nanopore reads from the Kraken2 step were re-analyzed for each sample. Unclassified reads (Q ≥ 10; Dorado sup) were length-filtered for any reads below 3,000 base pairs (Tables S8-9). The PhaBOX2 in end-to-end mode was run with its integrated modules in a single pass against the following bundled databases synchronized to the International Committee on Taxonomy of Viruses (ICTV) 2024 release (34), for further determining virus identification (PhaMer), lifestyle prediction (PhaTYP), taxonomic classification (PhaGCN), host prediction (CHERRY), and protein annotation (PhaVIP). The standalone implementation mirrors the PhaBOX web server while enabling batch processing on large metagenomes; the PhaBOX release (v2.1), database release (phabox_dv_v2_1), and command line were recorded in the run log (Figure S6).

We further analyzed the subset of previously unclassified reads predicted as viral that exceeded 40,000 bps and met a medium confidence threshold (Table S9). For these long viral contigs, we conducted completeness assessment, phenotype annotation (lifestyle prediction and host assignment), and open reading frame (ORF) prediction using PhageScope, an integrated phage-analysis pipeline based on recent bioinformatics tools with curated bacteriophage databases (49).

## Declarations

### Ethics approval and consent to participate

Not Applicable

### Consent for publication

Not Applicable

### Availability of data and materials

All codes related to this project are available under an open-source license at https://github.com/hshimlab. For nanopore data acquisition, we used MinKNOW v23.11.2 and MinKNOW core v5.8.0. For rapid nanopore data analysis, we used EPI2ME Labs v23.11.2 and Dorado v7.2.11. For taxonomic and AMR analysis, we used Kraken2 v2.1.3 and Abricate v1.0.1. For phage analysis, we used PhaBOX v2.1 and PhageScope. For data analysis and visualization, Python v3.6.4 and R v4.5.1 were used.

### Authors’ contributions

Experiments were primarily conducted by A.M. and H.S. Chemical and microbial analyses were led by A.M. and D.K., pathogen analyses were performed by B.L. and B.Q, and AMR and phage analyses were conducted by H.S. The study was conceived by H.S., and all authors contributed to writing the manuscript.

## Acknowledgments

We thank Cindy Drake at Pacific Agronomics for compost analysis, and Vince Roos and Robert Willmott at the Jordan College of Agricultural Sciences and Technology of California State University, Fresno, for sampling logistics.

